# Phylodynamic analysis of an emergent *Mycobacterium bovis* outbreak in an area with no previously known wildlife infections

**DOI:** 10.1101/2020.11.12.379297

**Authors:** Gianluigi Rossi, Joseph Crispell, Tanis Brough, Samantha J. Lycett, Piran C. L. White, Adrian Allen, Richard J. Ellis, Stephen V. Gordon, Roland Harwood, Eleftheria Palkopoulou, Eleanor L. Presho, Robin Skuce, Graham C. Smith, Rowland R. Kao

## Abstract

Understanding how an emergent pathogen successfully establishes itself and persists in a previously unaffected population is a crucial problem in disease ecology. In multi-host pathogen systems this problem is particularly difficult, as the importance of each host species to transmission is often poorly characterised, and the epidemiology of the disease is complex. Opportunities to observe and analyse such emergent scenarios are few.

Here, we exploit a unique dataset combining densely-collected data on the epidemiological and evolutionary characteristics of an outbreak of *Mycobacterium bovis* (*M. bovis,* the causative agent of bovine tuberculosis, bTB) in a population of cattle and badgers in an area considered low-risk for bTB, that has no previous record of either persistent infection in cattle, or of any infection in wildlife.

We analyse the outbreak dynamics using a combination of mathematical modelling, machine learning and Bayesian evolutionary analyses. Comparison to *M. bovis* whole-genome sequences from Northern Ireland confirmed this to be a single introduction of the pathogen from the latter region, with evolutionary analysis supporting an introduction directly into the local cattle population at least six years prior to its first discovery in badgers. Once introduced, the evidence supports *M. bovis* epidemiological dynamics passing through two phases, the first dominated by cattle-to-cattle transmission before becoming established in the local badger population.

These findings emphasise the importance of disease surveillance for early containment of outbreaks, in particular for pathogens not causing immediately evident symptoms in the infected host, and highlight the utility of combining dynamic modelling and phylogenetic analyses for understanding the often complex infection dynamics associated with emergent outbreaks.

## 1. Introduction

Pathogens able to spread at the interfaces between livestock, wildlife, and humans are one of the most serious threats to human health, wildlife conservation, and livestock economic sustainability [1,2]. Generally, the spread of a pathogen is enhanced when it co-circulates in multiple sympatric host species, as interspecific and intraspecific transmissions can complement each other, resulting in pathogen persistence [3,4].

*M. bovis*, a member of the *Mycobacterium tuberculosis* complex (MTBC) [5], is responsible for bovine (or animal) tuberculosis (bTB) in domestic cattle and a range of wild mammals [6], including European badgers and deer in Great Britain and Ireland [7–9], deer and wild boar in the Iberian Peninsula [10,11], deer and elk in Michigan, US [12], possums in New Zealand [13,14], and water buffalo in South Africa [15].

In Great Britain and Ireland several studies have established an association between the presence of infected badger populations and the persistence of bTB in cattle [16–18]. More recently, researchers have been able to demonstrate that the same *M. bovis* strains are co-circulating in domestic cattle and sympatric badgers in endemic bTB areas, first using the pathogen’s DNA genotyping techniques [19,20], and later using whole-genome sequencing [21,22]. Despite the efforts made by governments to control and eradicate bTB, the last decades have seen an increase in the number of cases and substantial expansion of bTB endemic areas, in particular in England and Wales [5,23]. The eradication efforts of this disease in England alone, costs the UK government around 100 million pounds per year [8,24,25].

Collecting reliable and up-to-date data about wildlife populations can present many challenges at broader scales [26] and while over the years many studies focused on characterising specific badger populations (see [27], and references therein), the lack of information in some areas might prevent the design of effective disease control practices when bTB is introduced. In addition, broad surveys across England have shown that, since the mid-1980s, the estimated number of badger social groups has been increasing by 2.6% annually, contributing to the uncertainty around their level of contributions potential as bTB reservoir in different regions [28]. However, reliable estimates of badger density, movement, and potentially infectious contact patterns are poorly recorded in most regions of Great Britain and Ireland, with only few populations such as at Woodchester Park (Gloucestershire, England) subject to denser sampling, in this case since the 1980s [29].

Further complications arise from the difficulty of estimating the true prevalence of *M. bovis* infection in badgers, as well as in domestic cattle. First is the elusive nature of *M. bovis*: the bacillus is characterised by slow replication with the potential for latent periods of variable length within the host [30,31]. In addition, the accuracy of currently available diagnostic tests is suboptimal in both cattle [32] and badgers [33]. These factors contribute to obscuring the relative roles of the two species in bTB maintenance and spread, and hampering the control and surveillance strategies. In particular, if both species are able to maintain the disease, control efforts focused on only one will be ineffective in achieving eradication [22].

One of the main goals of the current control and eradication strategy in GB is to prevent bTB from becoming established in new non-endemic areas, in particular those officially categorised as “low-risk” [34]. Within the Low Risk Area of England (LRA), the eastern part of the Cumbria county in north-west England (hereafter referred to as ‘East Cumbria’), has recently experienced a bTB outbreak of unusual magnitude and duration for this area. The outbreak began in 2014 and by mid-2019, through enhanced TB surveillance testing of all cattle herds in the affected area of East Cumbria, it had resulted in the detection of 39 breakdowns (positive cattle herds) across 33 premises [35]. The outbreak was caused by a strain of *M. bovis* (genotype) new to England, but previously observed in Northern Ireland, which was shown by preliminary molecular analyses to be the likely area of origin (Skuce, personal communication). Surveillance of ‘found dead’ wildlife (badgers and wild deer) for *M. bovis* infection was initiated in September 2016 in the area by the Animal and Plant Health Agency (APHA) [35]. By August 2018 three (out of 52 inspected) roadkill badgers had been found to be infected, all of them with the same bacterium genotype previously isolated from local cattle herds. As a result of this epidemiological link, a badger cull was licensed in a defined area within the affected part of East Cumbria with the aim of eradicating the disease in badgers and cattle. During the first season of culling operations in the autumn of 2018, 11% of all the removed badgers were found to be infected, all animals with the same genotype, suggesting that *M. bovis* infection has become established in the local badger population.

The aim of this study was to shed light on the dynamic spread of *M. bovis* when introduced in a two-host system in a non-endemic area, for which the outbreak in East Cumbria provided us with a unique opportunity to closely study. Here, we describe the East Cumbria outbreak’s spatial and temporal characteristics, and identify the factors which led *M. bovis* infection becoming established in a wildlife population, while estimating the number of intraspecies and interspecies transmission events. Our approach includes the use of forensic molecular epidemiology [36], since over 60 isolates of *M. bovis* from the outbreak with usable whole genomic sequences were available at the time of writing; complete with precise metadata including dates, locations and, for cattle, animal and farm identifications.

Results of this study are an important step toward a deeper understanding of bTB introduction and establishment into non-endemic areas, thus assisting with the process of disease risk assignment and future policy decision making by the animal health authorities.

## 2. Results

### 2.1. Outbreak description

In November 2014, typical bTB lesions were detected via routine slaughterhouse inspection of a seven month-old male calf from a dairy herd in East Cumbria. Bacteriological culture of the lesions yielded an unusual genotype of *M. bovis*, designated 17:z by APHA (Figure 1, A). Following this first report, 23 more cattle were confirmed to be infected with the same strain in East Cumbria, with the last of these detected in November 2018 (at the time of writing). Further cattle were declared bTB positive (using the tuberculin skin test and/or supplementary interferon-gamma blood tests) during this period in the outbreak area, although an *M. bovis* bacilli could not be isolated. The 24 cattle infected with *M. bovis* 17:z genotype included three animals detected outside the outbreak area but still in Cumbria, as well as three in the neighbouring counties of Lancashire (two) and Yorkshire (one), all deemed likely to be part of the same outbreak due to associations through contact tracing. The index animal in this outbreak was a homebred calf that had never left its birth farm until it was moved to slaughter. Therefore, this animal could not have been the “case zero” of this outbreak. We attempted to trace back the first infected individual introduced in the outbreak area by analysing all the Cattle Trace System dataset records that included animals born in Northern Ireland or in the Republic of Ireland from 2009 to 2014. Tracing back the direct movements from Northern Ireland to the outbreak area indicated a limited number of “first arrival” premises (on average 9.3 per year, range 5-15), but unfortunately it did not reveal an obvious first introduction. Conversely, searching for indirect links between Northern Irish farms by selecting other British farms with links to the Cumbrian outbreak which previously imported animals from Northern Ireland, provided too many potential “arrival” premises (on average 216.5 and range 165-247, farms in the outbreak area per year).

**Figure 1.**
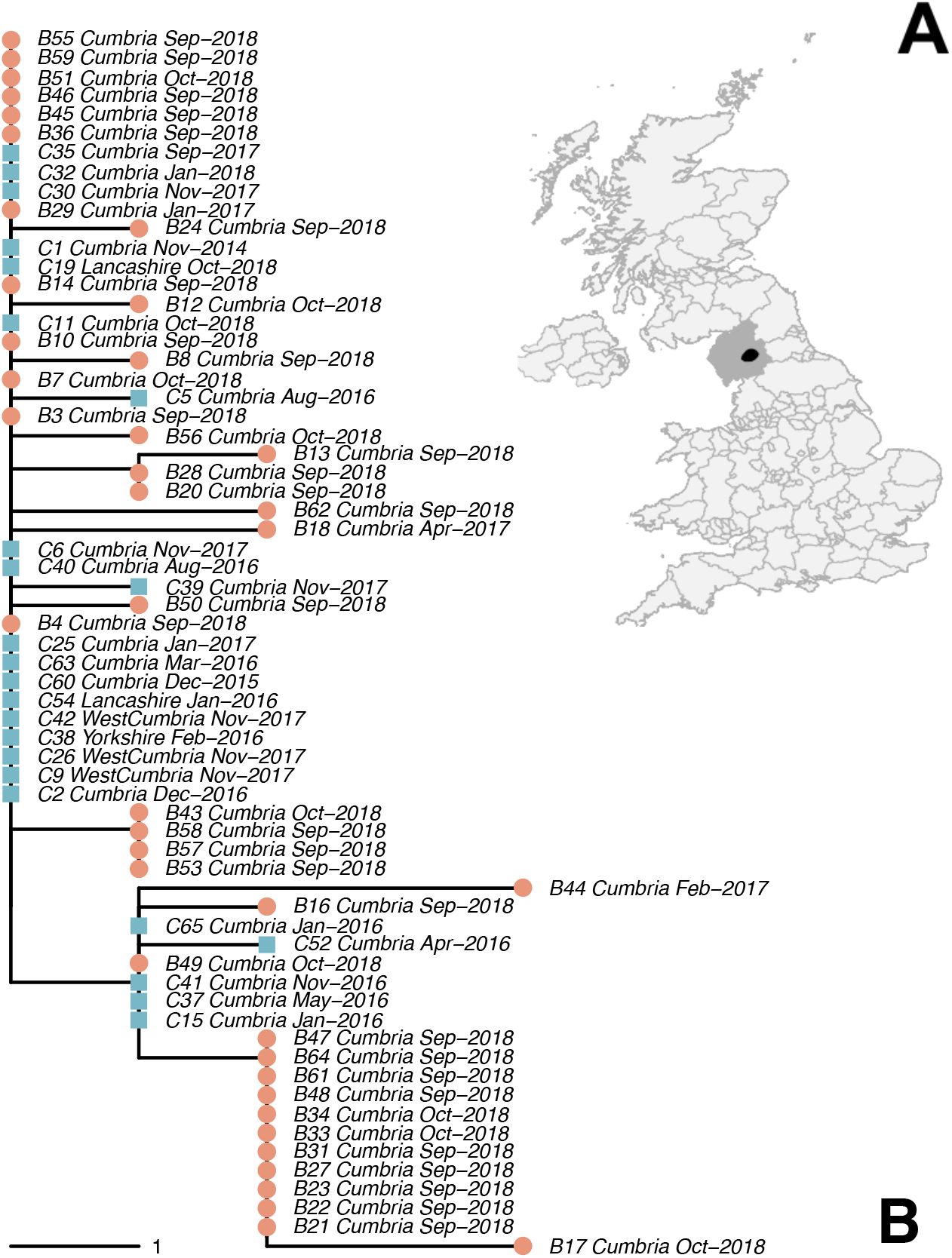
East Cumbria outbreak area and phylogenetic tree. A: East Cumbria outbreak area (black) location in Cumbria (dark grey). B: *M. bovis* genomes phylogenetic tree (distance calculated in Single Nucleotide Variants, SNVs). Red dots represent badgers, and blue square cattle, and the label report the code assigned to each individual, sampling location and sample date.

During 2016 and 2017, respectively, two and 35 roadkill badger carcasses were reported to the local authorities within the designated outbreak area of East Cumbria. Three of the badger carcasses, retrieved respectively in January, February, and April 2017, were positive for *M. bovis* on culture (while two carcasses were unsuitable for inspection) (Figure S1.1), and all three positive animals were infected with the 17:z genotype.

The identification of infected badgers led Defra, following a public consultation, to issue a badger culling licence in a specified section of the outbreak area in the autumn of 2018. Culling operations from September to November 2018 resulted in 602 culled badgers, of which 369 were submitted for post mortem inspection and laboratory testing (Defra 2019). In total, 42 were culture positive for *M. bovis* and of those 38 isolates yielded a whole genome sequence of sufficient quality to enable phylogenetic analysis [37].

Data on found-dead surveillance of 2018 and 2019, but prior to the second culling season, included an additional 42 retrieved carcasses (29 and 13, respectively), all negative to *M. bovis*, except for 15 that were unsuitable for post-mortem inspection, and three still pending at the time of data gathering.

### 2.2. Outbreak phylogeny and Northern Ireland isolates

The phylodynamic tree of the East Cumbria outbreak is reported in Figure 1 (B). Early analyses identified a genotype usually found in Northern Ireland; further evidence showed the existence of cattle movements from this area to England and Wales. Coincidentally, the origin area included the recently completed Test, Vaccinate or Remove (TVR) trial area [38] in Northern Ireland (Skuce, personal communication) where extensive *M. bovis* whole genome sequencing had already been done – these isolates were included in the current analyses.

The complete phylogenetic tree (Figure 2) confirmed the association between the *M. bovis* circulating in the Northern Irish TVR area and in East Cumbria; thus it appeared that the East Cumbrian outbreak likely originated from the dominant strain circulating in or around the TVR area (Figure 2, orange branches) imported by movement of infected cattle, though the first introduction was not identified.

**Figure 2.**
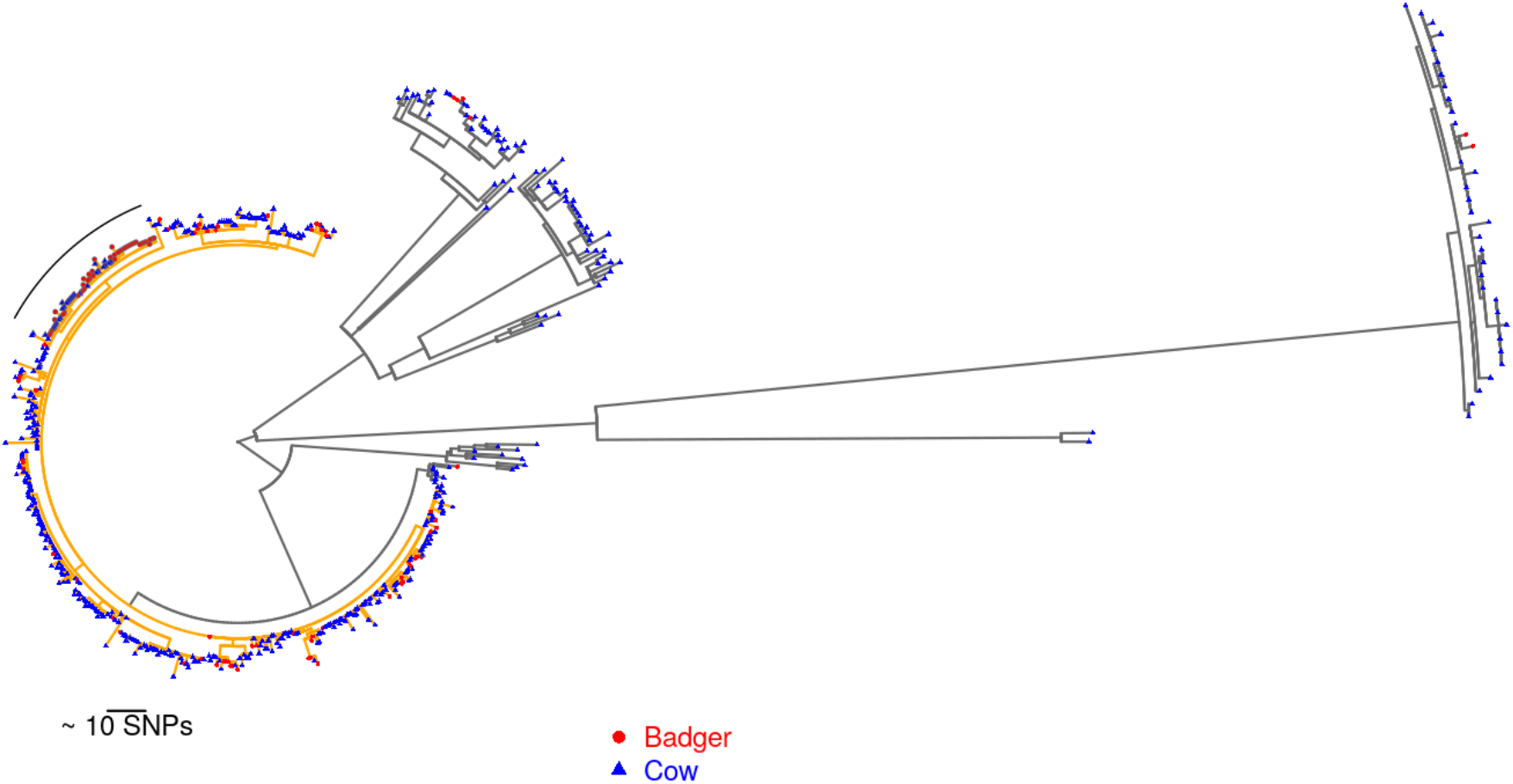
East Cumbria and TVR *M. bovis* phylogenies. A maximum likelihood phylogeny of the *M. bovis* genomes sourced from Cumbria and the Test, Vaccinate, or Release (TVR) area in Northern Ireland. The tree is rooted with the *M. bovis* reference genome AF2122/97. The *M. bovis* genomes sourced from infected cattle and badgers in Cumbria are highlighted with a black semi-circle at the top left. The branches of the clade containing the *M. bovis* genomes sourced from Cumbria, and those from the TVR area that are most similar is highlighted in orange.

### 2.3. Epidemiological signatures in genetic data

Following our previous approach [22], epidemiological signatures in the sequence distances were identified using Boosted Regression Trees (BRT) [39], which combines decision trees and boosting techniques [40].

As previously, the dependent variable was the genetic distance between *M. bovis* strains, expressed as single nucleotide variants (SNVs); for explanatory variables we calculated 18 relational covariates for each pair of sampled animals. These covariates are listed in Table 1 and are divided into four categories: temporal (1 covariate), spatial (2 covariates), group (3 covariates), and contact networks (12 covariates).

**Table 1.** Epidemiological covariates tested in the BRT algorithm. List of epidemiological covariates tested in the boosted regression trees (BRT) algorithm against the isolates genetic distance, calculated in number of SNVs. The four columns indicate in which model the covariate has been used.

| Category | Covariates | Description | All samples model | Cattle model | Badger model | Inter-species model |
| --- | --- | --- | --- | --- | --- | --- |
| Temporal | SampleTimeDist | Time distance (in days) between the isolates sample dates | YES | YES | YES | YES |
| Spatial | SampleGeoDist | Geographical Euclidean distance between the isolates sampling locations | YES | YES | YES | YES |
| Spatial | ParcelsDist | Minimum geographical Euclidean distance between the land parcels where isolates have been sampled | YES | YES | YES | YES |
| Group | SameSpecies | Isolates from same species animals (YES/NO) | YES | NO | NO | NO |
| Group | SameGroup | Isolates from same herd or sett animals (YES/NO) | YES | YES | YES | NO |
| Group | Sizes | Size coefficient (squared root of product) of the isolates herd/social groups | YES | YES | YES | YES |
| Network | ShortestPathSNet | Shortest path length between isolates nodes in single-species network | YES | YES | YES | NO |
| Network | InfNodesInPathSNet | Number of infected nodes in shortest path between isolates in single-species network | YES | YES | YES | NO |
| Network | DegreeSNet | Degree coefficient (squared root of product) of isolates nodes in single-species network | YES | YES | YES | NO |
| Network | SameCommSNet | Isolates' nodes in same community in single-species network (YES/NO) | YES | YES | YES | NO |
| Network | ShortestPathLPNet | Shortest path length between isolates nodes in spatial (land parcels) network | YES | YES | YES | YES |
| Network | InfParcInPathLPNet | Number of infected nodes in shortest path between isolates in spatial network | YES | YES | YES | YES |
| Network | DegreeLPNet | Degree coefficient (squared root of product) of isolates nodes in spatial network | YES | YES | YES | YES |
| Network | SameCommLPNet | Isolates' nodes in same community in spatial network (YES/NO) | YES | YES | YES | YES |
| Network | ShortestPathML | Shortest path length between isolates nodes in multi-layer network | YES | YES | YES | YES |
| Network | InfNodesInPathML | Number of infected nodes in shortest path between isolates in multi-layer network | YES | YES | YES | YES |
| Network | DegreeML | Degree coefficient (squared root of product) of isolates nodes in multi-layer network | YES | YES | YES | YES |
| Network | SameCommML | Isolates' nodes in same community in multi-layer network (YES/NO) | YES | YES | YES | YES |

The BRT model on the full dataset (1,711 observations) was able to explain 41% (pseudo R^2^) of the variability on the test dataset, while the Root Mean Squared Error (RMSE) was 0.94. To test the robustness of this model we ran the same analysis on the same dataset but with an increasing percentage of randomly reassigned values for the dependent variable (SNV distance) observations. The results (Figure S3.1) showed that even for a limited percentage of re-assigned data (10%) the model underperformed substantially, explaining only 28% of the variation on average. The full model results (Figure 3) showed that the most important covariate to explain the SNV distance between isolates was the time between isolate samplings (25.2%), followed by the hosts populations’ size (14.3%) and the geographical distance (considering both isolate sampling locations and between isolates land parcels, respectively 11.3 and 9.3%). Among the contact network covariates, the degree in multi-layer (9.2%) and land parcels networks (9.1%) were the most important, while the covariates related to the single-species networks were not important (all lower than 4%). Partial dependency plots are reported in Figure S3.1.

**Figure 3.**
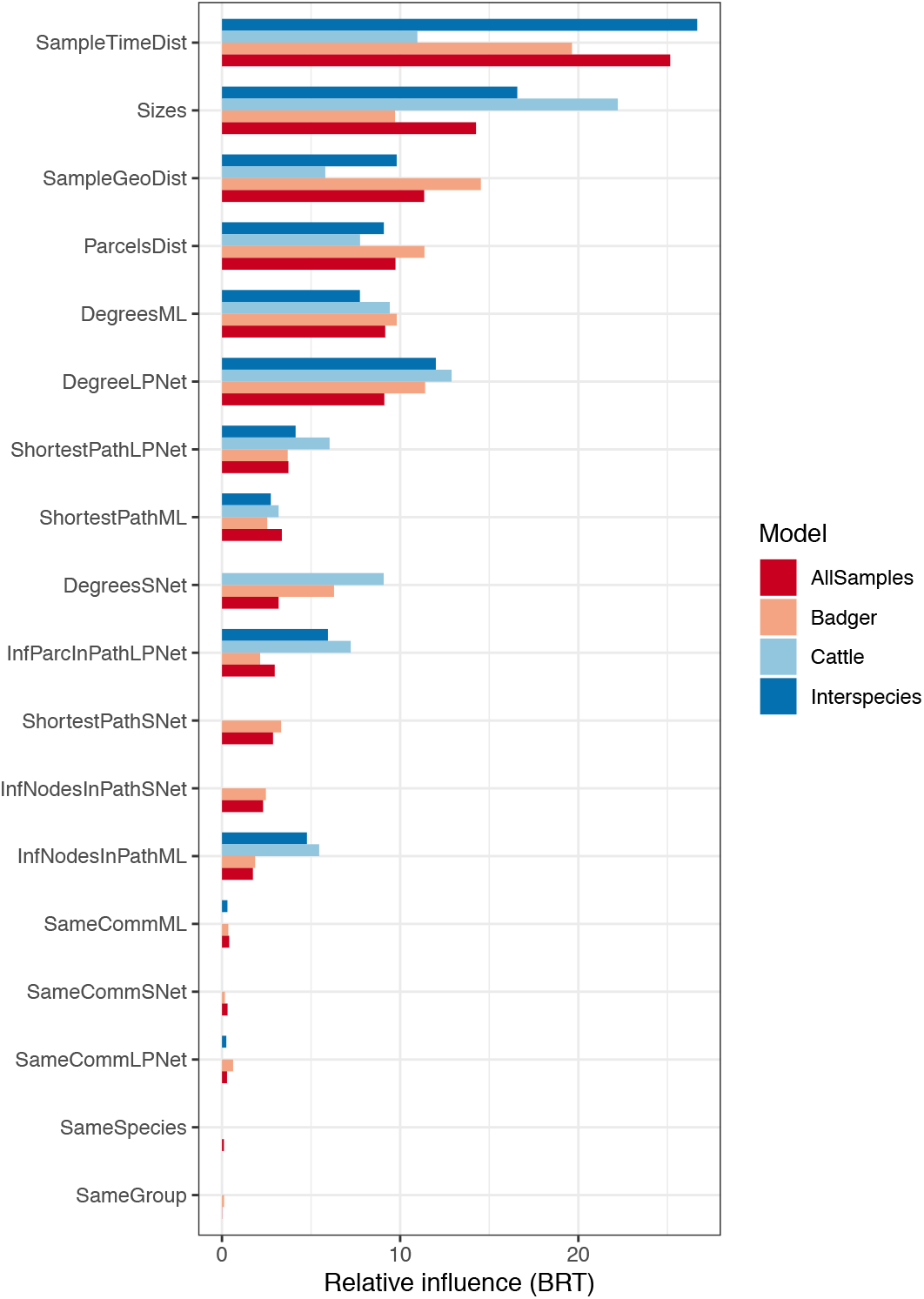
Covariate influence in the BRT models. The relative influence of the 18 epidemiological covariates calculated by the Boosted Regression Trees (BRT) model. Bars colours correspond to the four models run with different sub-samples of the dataset: dark-red for the full model account all samples, light red for the badger-to-badger model, light-blue for the cattle-to-cattle only model, and dark-blue for the interspecies (badger-to-cattle) model.

We ran a further three BRT models: two of them considered the within-species interactions only, cattle-to-cattle (153 observations), and badger-to-badger (820 observations). Conversely, the third model considered cross-species interactions (738 observations). These models were able to explain respectively 16%, 43% and 36% of the variation, while the respective RMSE values were 0.82, 1.04, and 0.96.

The covariates influence rankings in the badger and inter-species models were similar to the full model (sampling time distance the most influent, respectively, 19.6 and 26.7%), with the exception that in both cases the degree in the land parcels network was the third most influential covariate (11.4 and 12.0%). In the badger model the population size dropped to sixth (9.7%), while the geographic distance between sampling locations was second (14.5%). The cattle model showed the most differences with the full model, with group size and degree in the land-parcel network being the most influential covariates (22.2 and 12.9%), while sampling time distance was third (11.0%).

### 2.4. Pairwise transmission probability and most likely transmission tree

The pairwise transmission probability was calculated using the Kolmogorov Forward Equations (KFEs)[41], and the pairwise transmission probability matrix was reported in Figure 4. The KFEs methodology was used to calculate the probability of observing a pair of bTB infected hosts given their sampling time and the genetic divergence between their bacterial isolates, assuming that a direct transmission occurred between the pair.

**Figure 4.**
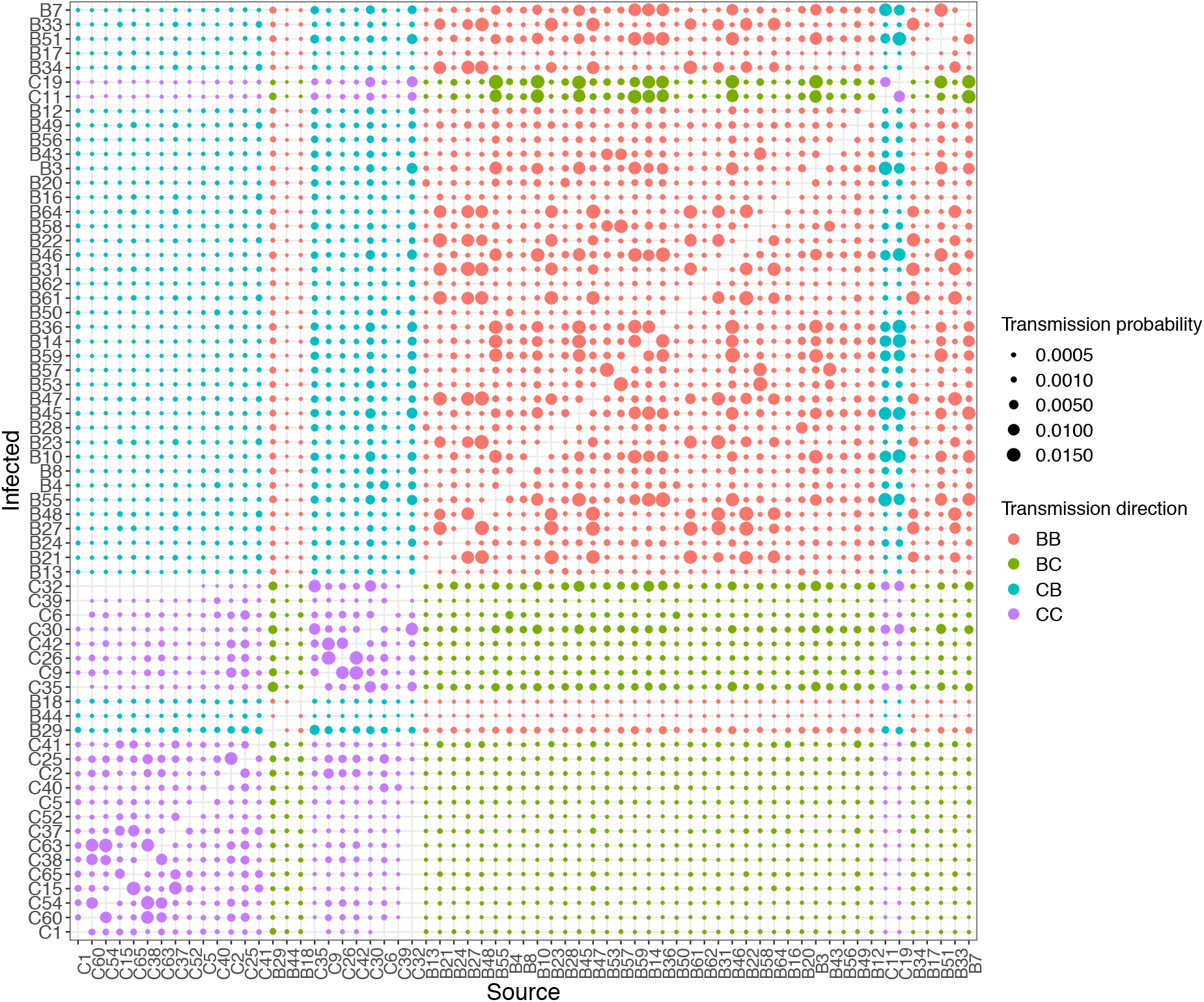
Transmission matrix. Pairwise transmission probabilities between infected animals in the East Cumbria outbreak. Animals are reported in *x* (source animal) and *y* (infected animal) axes in the order they have been sampled, and they are labelled from one to 65 and the species name (B for badgers and C for Cattle). Different colours correspond to transmission directions (red: badger-to-badger, green: badger-to-cow, light-blue: cattle-to-badger, and magenta: cattle-to-cattle).

The animal that infected the first detected cattle (C1) with *M. bovis* might have escaped detection, since the transmission probabilities from other sampled animals are low (median 0.13 × 10^−3^, range 0 – 1.98 × 10^−3^). Similarly, two out of three roadkill badgers (B44 and B18) had, respectively, the lowest and the third lowest average and maximum transmission probability from other sampled animals (see Figure 5). This indicates that animals infected early in the outbreak likely escaped detection (either the 4-year herd testing or carcass inspection), in particular the “case zero” (i.e. the first cow imported infected with the 17:z genotype of *M. bovis*).

**Figure 5.**
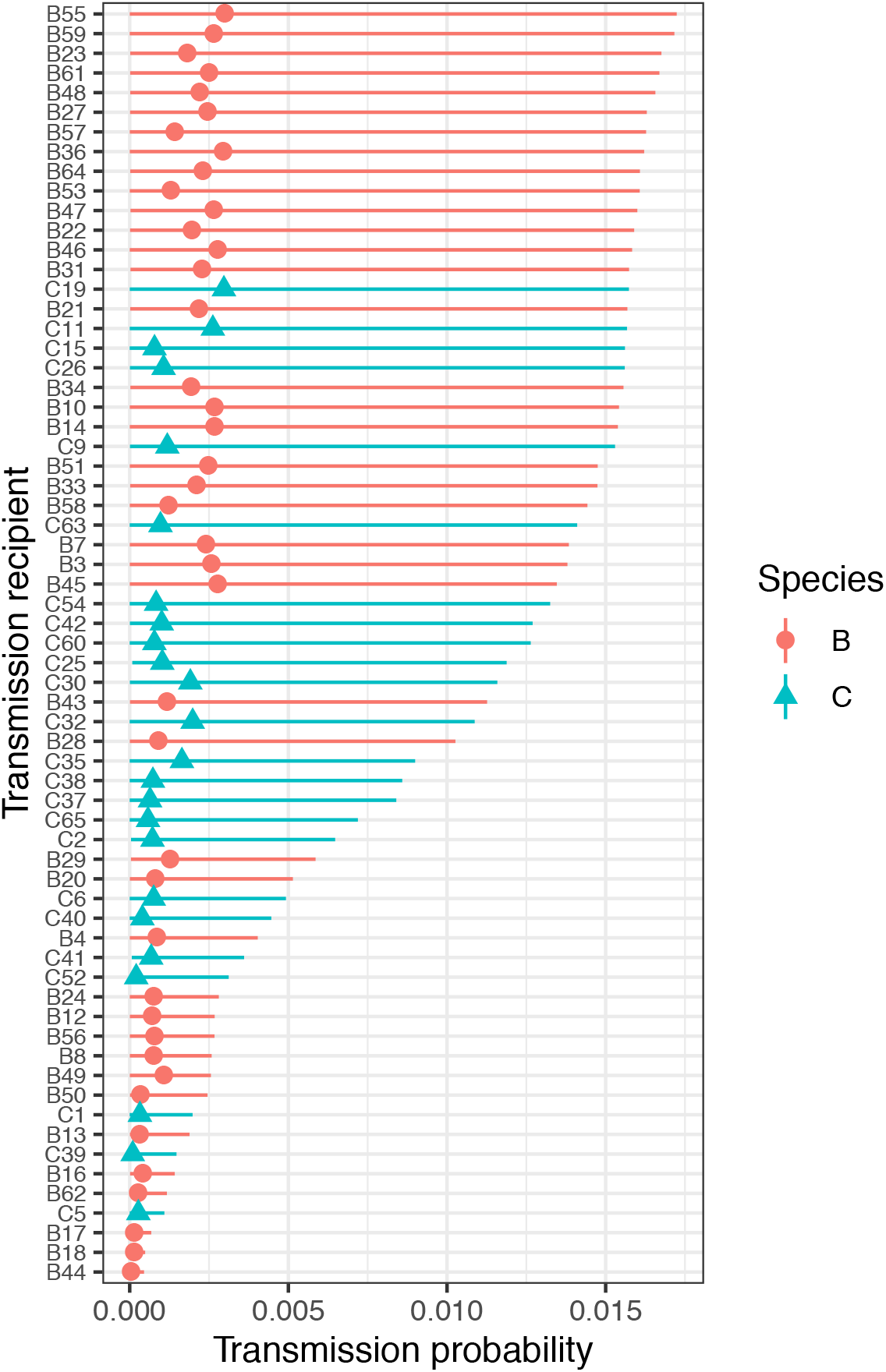
Transmission probability to sequenced animals. Average (dots/triangles) and range (line) of infection probabilities (x axis) from all sampled infection sources to each individual animal (y axis). Red lines/dots correspond to badgers and blue lines/triangles to cattle.

**Figure 6.**
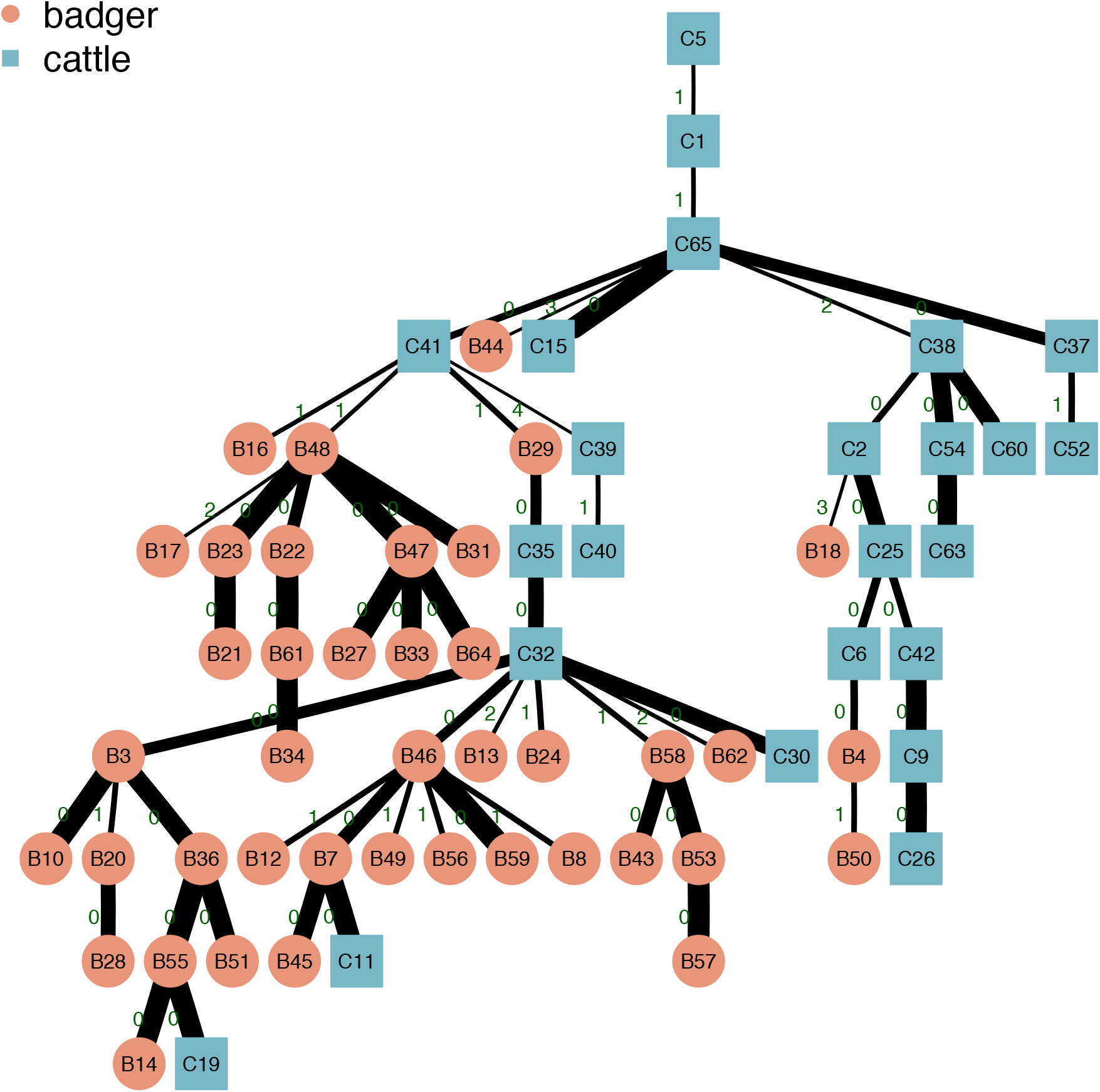
East Cumbria *M. bovis* outbreak transmission tree. Most likely transmission for the sampled infectious individuals in the East Cumbria outbreak (transmissions from top to bottom). Red circles correspond to badgers, and blue squares to cattle. Edge thickness is proportional to the pair transmission probability and the edge label (dark green) indicates the SNV distance between the two individuals.

In general, within-species transmission probabilities were higher than between-species ones (Figure S4.1), and the phylogenetic root-to-tip temporal signal was strong as well (R^2^ = 0.39, p-value ~0; see Figure S4.2). This was consistent with the BRT model results, where the temporal signal was identified as the most important factor to predict the SNV distance. We computed 10,000 “random trees” built by selecting random transmissions (except the ones for which the probability was zero because the cattle’s lifespans did not overlap). Results showed that random trees had a median [95^th^ quantile] of 17[13–21] cattle-to-cattle, 26[21–31] badger-to-badger, 14[10–19] cattle-to-badger, and 6[2–10] badger-to-cattle transmissions (Figure S4.3). We computed the most likely transmission tree by selecting the transmissions with highest probability within the sampled animals while avoiding loops in the tree (see [41]). The best tree (Figure 4) showed that most of the transmissions in this system likely happened within-species, i.e. 20 cattle-to-cattle and 29 badger-to-badger, respectively. Conversely, inter-species transmissions comprised 12 cattle-to-badgers incidents, and 3 badgers-to-cattle. When comparing these results with the randomly computed trees, the most-likely tree showed a lower number of cross-species transmissions and a higher number of within-species transmissions, although the estimates fell in the 95^th^ quantile of the random trees ones.

### 2.5. BASTA analysis

The transmission rates estimated by Bayesian Structured coalescent Approximation (BASTA) [42] on the 10 different sub-samples suggest that transmission from cattle-to-badgers occurred much more frequently (at least an order of magnitude) than transmission from badgers-to-cattle in Cumbria (Figure 7). In addition, there is little support for the inclusion of badgers-to-cattle transmission in the structured population model. In contrast, there is strong support for transmission of *M. bovis* from the sampled Northern Ireland area into the Cumbria area via the cattle population. Figure S5.1 shows the rate estimates that BASTA produced when no genomic data was provided, therefore only sampling dates were used. These analyses were conducted to determine how much signal there was in the genomic data to support the transmission rates being estimated. Given the contrasting rates shown in Figure 7 and Figure S5.1, there was strong evidence that there was sufficient signal in the *M. bovis* genomic data to estimate the transmission rates. Lastly, there is good agreement across the 10 sub-samples, suggesting that the estimated rates were robust to any inter-sub-sample variation.

**Figure 7.**
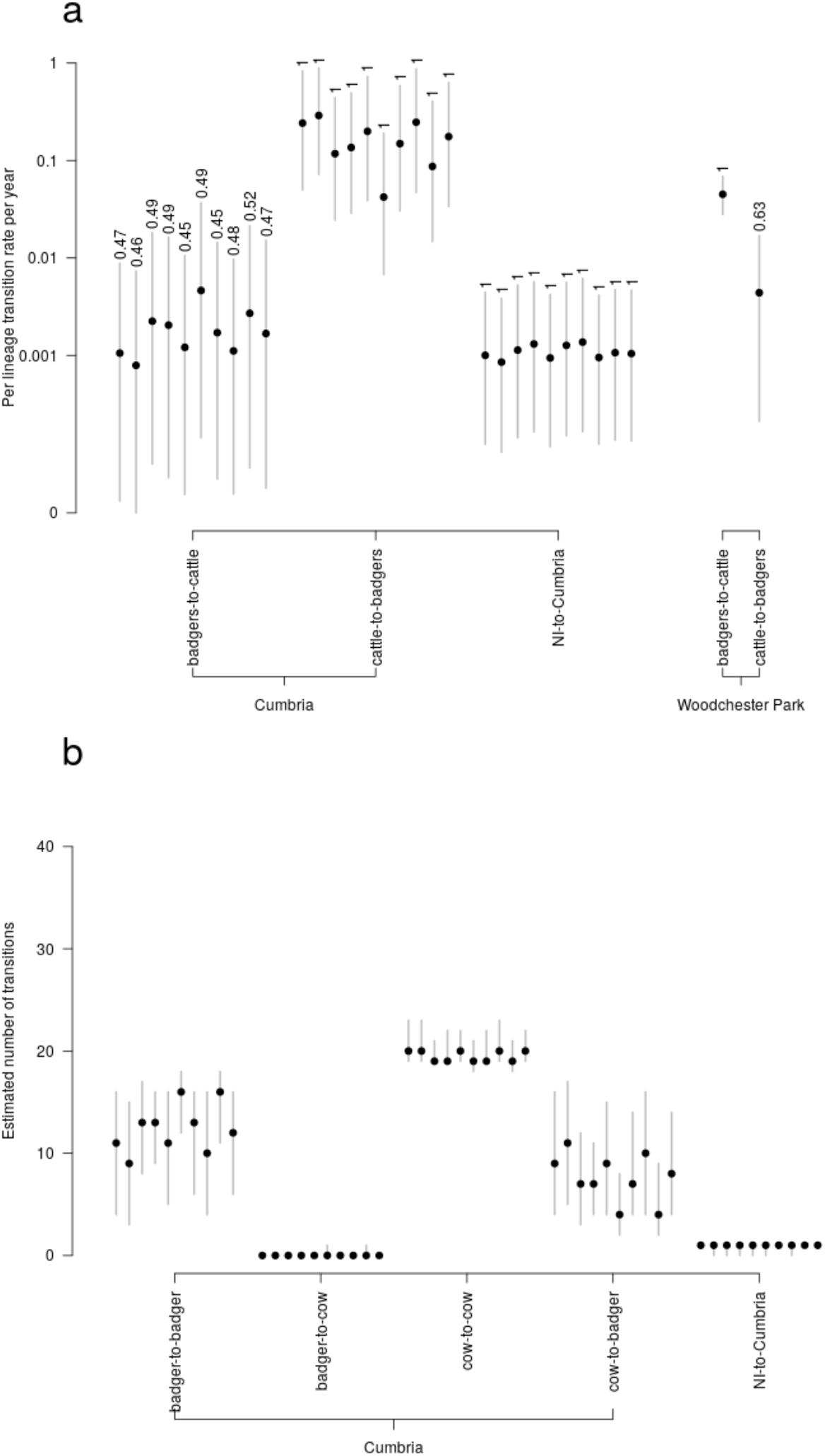
Transmission rates and number of transmissions estimated with BASTA. a) Inter-species and Northern Ireland to Cumbria *M. bovis* transmission rates (y axes, log scale) estimated using BASTA, based on analyses that used the genomic data and compared with the results for Woodchester Park [22]; b) estimates from BASTA of the number of transmission events (y axes) between the sampled cattle and badgers.

Analyses in BASTA leveraging the temporal signal in the *M. bovis* genomic data were used to estimate the timing of *M. bovis* transmission from the cattle in the TVR area to cattle in Cumbria (Figure S4.4). Whilst credible intervals around these estimates were very broad, the transmission event was estimated to have occurred in March 2011 (lower 2.5% bound estimate: August 2001; upper 97.5% bound estimate: April 2014). This reflect the slow and variable replication rate characteristic of *M. bovis*.

## 3. Discussion

While it is accepted that cattle movements are responsible for transmission on a wide spatial scale [23,43,44], uncertainty remains around the relative roles of cattle and badger in maintaining and spreading *M. bovis* infection at local scales, complicating the formulation and execution of control policies. Our analyses of the East Cumbria outbreak highlight this dynamic, with the introduction of *M. bovis* most likely being caused by a cattle movement from Northern Ireland, followed by a more complex spread among local cattle and badgers. Identifying the source of infection for the index case in cattle in this outbreak (marked as C1 in the figures) would be a crucial piece of information. Should a badger be the most likely infection source then this would imply an earlier establishment of the disease in wildlife (i.e. before or during 2014). Our results suggest another infected bovid as the most likely bTB source for the index case, but the transmission from known cases is poorly supported as the transmission probabilities from the other sampled animals to the index are generally low (Figure 5).

Overall, both the likely transmission tree selection (obtained with the KFEs) and the BASTA analysis indicate that most transmissions happened within-species, and that cattle-to-badger transmission has played a more important role than badger-to-cattle transmission, with similar outcomes obtained using two different analyses.

Our hypothesis, which is supported by the described results, is that after an initial seeding of *M. bovis* into the local East Cumbria cattle population, the infection subsequently became established in local badgers. Once that happened, rapid spread of the pathogen led to the observed 11% prevalence in the autumn of 2018, with peaks up to 20.9% in the core area [37]. Given the historically low prevalence of *M. bovis* in this area [45], the immunological naivete of this specific badgers population might have facilitated a quicker spread, although more research in this direction is needed. From a disease dynamic point of view, the relatively higher number of predicted cattle-to-badger transmission events suggests that the establishment in badgers may happen when infection pressure from the sympatric cattle population reaches a threshold, rather than a single transmission event.

A concern might be that transmission tree outcomes have been affected by the sampling timeline, since early in the outbreak there are less *M. bovis* sampled sequences and most come from cattle, while later on the majority of sampled isolates come from badgers. While the BASTA analysis is designed to specifically reduce the effects of unbalanced sampling [42], our conclusions are also supported by a low genetic diversity in the recovered badger *M. bovis* isolates, which points to a relatively recent outbreak in this population. Nonetheless, the BASTA analysis was able to exploit this low genetic diversity, as it is shown by the different results of this model when neglecting the genetic information.

Similar to a previous study in a separate population [22], the Boosted Regression Trees (BRT) analysis indicated temporal bacterial isolation differences, and spatial distance between hosts, as the most important factors to predict inter-*M. bovis* genetic distance (SNVs). Another important variable was host population sizes, with bigger groups (farms or setts) linked to higher SNV distances. This result was particularly strong for the BRT models which included cattle isolates, and it is consistent with farms size being a risk factor for bTB [46,47].

Including different contact networks metrics variables in the BRT model can help to understand which transmission mechanism is more relevant in this system. Among these variables, the number of adjacent land parcels, corresponding to the spatial network’s degree, was equal (full model) or more important (cattle, badger and inter-species models) than other metrics. This points to fine-scale spatial effects which cannot be entirely explained by the simple distance between samples locations. On the contrary, the single-species network metrics (animal movements for cattle and sett adjacency for badgers) were not significant in any models. This might be the result of complex interactions between the two species that cannot be explained when considered individually, as previously suggested [48]. However, these interactions become evident when both species contact networks are included in a single framework such as multi-layer network [49].

The badger-to-badger model was able to predict the SNV distances in the test dataset slightly better than the full model (43% to 42%), despite reducing the sample size (from 1,711 observations in the full model, to 820 in the badger only model). Furthermore, we observed that the SNV distance variability explained by the cattle only model was comparable to that of the full model when 20% of the data were randomly shuffled. This was surprising since more data are collected on domestic animals and this should provide a better picture than the equivalent wildlife data. One potential explanation is that the selected variables for this model might better explain *M. bovis* dynamics in badgers than in cattle, at the local scale. However, when we consider this and the good explanatory power of the land parcels network metrics, we could speculate that the cattle dataset might hide some contact patterns. This could occur due to the inability to isolate *M. bovis* in all herd breakdowns or in all cattle, or due to unrecorded movements, with cattle grazing in several land parcels belonging to the same farm but not contiguous to the farm [50,51]. In general, the landscape and the farmland fragmentation and distribution in space, which is ignored when considering farms’ main building locations only, might play a crucial role in disease dissemination. This calls for surveillance and control strategies to be adapted to specific contexts and informed by detailed veterinary investigation and spatial information, such as land parcel distribution.

By comparing the East Cumbria outbreak with other bTB in other contexts leads to some important insights. In East Cumbria, badgers have played a lesser role in the local persistence of the disease, the converse to what observed in a bTB endemic area in Woodchester Park [22]. This may be due to the “age” of the outbreak. Whilst in newly infected areas, and therefore non-endemic, cattle-to-cattle and cattle-to-wildlife transmissions may dominate, in endemic areas the dynamic may have shifted towards a more complex dynamic, where wildlife can play a more important role. While in both cases breaking the transmission at the wildlife/livestock interface is critical, in outbreaks within low risk areas (non-endemic) it is crucial to prevent the establishment of bTB.

Questions remains as to how likely bTB is to get established in non-endemic areas, and how long it would take to detect it. From this perspective, the fact that the first introduction in this area (estimated to be 2011), and that the initial infected cattle were not detected until inspection at slaughter, indicates limitations of the bTB surveillance strategy in low risk areas and the importance of good biosecurity to reduce the risk of onward transmission to wildlife from introductions of cattle with undetected *M. bovis* infection. In order to mitigate this risk, in April 2016 Defra adopted mandatory post movement bTB testing of all cattle moved from high-risk to lower-risk areas [52].

To conclude, our analyses of the recent East Cumbria outbreak highlight how the transmission dynamics of *M. bovis* can change during the establishment in a non-endemic area, and how these changes affect the relative roles of wildlife and domestic animals in establishing and maintaining the infection. Our results suggest how genomic data and phylogenetic analyses are becoming fundamental to disentangle bacterial pathogen outbreaks, in particular when combined with epidemiological and network models [53]. Therefore, infrastructures for genomic surveillance can definitely help inform bTB and other endemic diseases control policies.

Finally, we highlight how local spatial dynamics might affect pathogen spread during the early phases of an outbreak. This makes the case for different diseases control strategies in endemic and non-endemic areas that take into account detailed characteristics about the system landscape and rearing practices, and for improving biosecurity, in order to achieve minimal transmission at the cattle/wildlife interface and therefore preventing the establishment of newly introduced diseases.

## Methods

### 4.1. Data description

#### 4.1.1. Sequences and metadata

Test positive cattle, found-dead badgers and most culled badgers in the Cumbria area were subject to post-mortem and culture of suitable tissues at the Animal and Plant Health Agency (APHA), with positive cultures subjected to genotyping and whole genome sequencing at the Central Sequencing Unit in Weybridge. 65 *M. bovis* whole-genome sequences were available from East Cumbria (65 in total, 24 from cattle, 3 from roadkill badgers, and 38 from culled badgers). The sampling timeline of the available sequences is reported in Figure S1.1. The metadata included a unique identifier, the sampling date, location coordinates (for badger isolates), and the farm’s county-parish-holding (CPH) code (for cattle sequences only). The isolates and raw sequence data were processed using the same pipeline as described by Crispell et al. [22].

The East Cumbria outbreak dataset included a further sequence from the same *M. bovis* genotype sampled in Scotland (Figure S1.2). The epidemiological investigation showed that the animal was imported from Northern Ireland for slaughter only, thus we did not consider it in the analyses.

#### 4.1.2. Badger population

A total of 160 badger setts were identified in and around the outbreak area in 2017/18 by the APHA, 117 of them were in the 2018 culling permit area. Badgers were both shot and trapped, with trapped animals subjected to post-mortem analysis, and population data (number of badgers removed, TB positive, negative, and TB status not determined) were available for 99 setts.

#### 4.1.3. Cattle population and outbreak area definition

To obtain all infected cattle life histories (movements, birth and death) we first matched the sequences’ unique identifier in the SAM dataset, then we extracted the data from the Cattle Tracing System (CTS) using the animal unique identifier.

The outbreak area was defined as the area within the minimum circle around the sequences sampling locations and all 160 badger setts, and adding to that all the parcels assigned to the infected farms which are contiguous to the above described circle (Figure 1, A). This study included all the farms active between 01/01/2010 and 31/12/2018 (which reported any cattle movements, births or deaths) directly located in the area or which owned a parcel in the outbreak area. The total number of selected cattle farms was 336.

#### 4.1.4. Northern Ireland TVR data

The Test, Vaccinate or Remove (TVR) trial in Northern Ireland ran from 2014 to 2018, and it was designed to determine whether a combination of vaccination and an animal side TB test could be an effective means of controlling *M. bovis* infection in badger populations. During this period *M. bovis* samples were taken from infected cattle and badgers in the area for culturing.

All positive cultures were sequenced at the Agri-Food and Biosciences Institute in Belfast (AFBI-NI). In addition, archived *M. bovis* isolates that were stored as part of routine surveillance operations in the TVR area prior to the start of the trial were selected for sequencing. These additional isolates were sourced from routine test and slaughter surveillance of cattle or road killed badgers.

From the TVR area in Northern Ireland, there were 544 *M. bovis* genomes sourced from infected cattle (479) and badgers (65), sampled from 1996 to 2017. The distribution of sampling times for the *M. bovis* genomes is shown in Figure S1.1.

### 4.2. Epidemiological information and machine learning analysis

An important part of this analysis involved investigating the population, temporal, spatial, and contact network signatures in the sampled *M. bovis* genomic data [22]. We computed the inter-sequence Single Nucleotide Variants (SNV) distance and then we fitted a Poisson regression model using the Boosted Regression Trees [39] model in R [54].

As mentioned above, we used population, temporal, spatial, and network covariates, which are listed in Table 1. Network covariates (shortest distance, same community, degree, and number of infected nodes in the shortest path) were calculated for three contact networks: single species, spatial, and multi-layer.

The *single species network* accounted for only the within-species potential contacts. The cattle population’s nodes corresponded to farms, and edges correspond to movements. In order to build these networks for each pair of cattle sequences, we only accounted for movements spanning from the previous year of the first of the two sequences sampled, to the year of the second one (i.e. if the two sequences were sampled respectively in 2015 and 2017, we computed the network by using movements recorded between 2014 and 2017). The badger population’s nodes corresponded to setts, and since most of the trapped badgers occurred near a sett, their sequences were assigned to the closest one. We used the Voronoi partition (or Dirichlet tessellation) to create the edges between setts, in order to avoid relying on an assumption for the distance at which two setts were connected. In this network configuration the inter-species sequences are considered not connected.

The *spatial network* was built by considering each land parcel as a single node. Two land parcels were considered connected by the proximity criterion: if they shared a border or if the borders were closer than 100 meters. In this case, the degree of a node in this network (i.e. land parcel) corresponds to the number of neighbouring parcels.

Finally, the *multi-layer network* accounted for all the previously described contacts combined. In this case, nodes are defined as land parcels as well, but on top of the proximity criterion, two land parcels could also be connected if they include two farms or setts which were connected in the single species network.

The network communities were defined using the Louvain method which optimises the modularity, as provided in the R package *igraph* [55].

The model was run on the complete dataset (All Samples, Table 1 and Figure 3), on the cattle and badger isolates only, and on the inter-species isolates. In all cases the train and test dataset were half of the observations, while in the cattle dataset we used 60% for training and 40% for testing, given the reduced observations compared to the others. We evaluated the models using the pseudo-R^2^ calculation comparing the observed and predicted SNV distances on the test datasets, and the Root Mean Squared Error (RMSE). These were both calculated using the package *caret* [56]. For BRT the relative influence of the covariates is determined by the times each variable is selected to split the data in a decision tree, which in turn is weighted by the improvement to the model fit that resulted from that variable being used at each split [39]. All models were fitted with a 10-fold cross validation.

Preliminary runs of the models were used to tune the BRT parameters in order to improve the predictions. These parameters were the learning rate, which controls the contribution of each tree to the final model, and the tree complexity, which corresponds to the number of nodes in the tree. For the full dataset model the learning rate was set to 0.005 and the tree complexity to 5. For the other model we set the learning rate to 0.003, 0.009, and 0.0075 for the cattle, badger and inter-species dataset, respectively and the tree complexity to 7, 1, and 10.

### 4.3. Pairwise KFE and transmission tree reconstruction

We calculated the transmission probability between pairs of animals where the *M. bovis* sequences have been sampled. We calculated this probability using the Kolmogorov-Forward Equations (KFE) methodology [41]. The KFEs consist of a set of ordinary differential equations which track the probability of a system to be in a given state through time [57–59]. In the pairwise transmission case, the system state was given by the combination of the two hosts disease progression state and, once an individual is infected, by the number of SNVs on its *M. bovis* strain. The underlying assumption was that, at the time of infection, the two pathogen strains found in the source and recipient hosts are identical. After the infection, the two strains start to replicate, and thus substitution on the pathogen DNA happens at a rate *μ* (*substitution rate*), generating SNVs. Because the two strains are diverging, we will call the SNVs found in a strain sampled in the source host A [recipient host B] as *divergent* SNVs, or *divA*[*divB*]. The sum of *divA* and *divB* results in the SNV distance between the two strains. In order to use this methodology, we have to provide three main pieces of information: the pathogens sequences, the two hosts sampling time, and an underlying model representing the disease progression. For bTB, we chose to use a simple Susceptible-Exposed-Infectious model [23,60], where susceptible can become exposed (or latent) after contact with an infectious host with *infection rate β*, and exposed hosts move to the infectious state with *transition rate σ* (or after a latency period of average *1/σ*). In this study, we provided the birth date of the cattle as well, in order to limit the time span where each cattle could have been infected first. For badgers we assumed a constant death rate, based on the observation that less than 0.1% of individuals would survive past 8-years of age [27]. The epidemiological parameters were chosen according the most recent literatures (see [41]), and in order to account for their variability, for each pairwise transmission we tested a 1’000 combination of randomly selected parameters combinations and chose the one returning the highest probability.

Following Rossi et al. [41], we assembled the most likely transmission tree by progressively selecting the pairs with the highest transmission probability, and excluding those not possible given the previously selected ones (e.g. if A⟶B and B⟶C are selected, B⟶A, C⟶B and C⟶A were going to be a priori excluded).

### 4.4. BASTA analysis

Given that the phylogenetic evidence suggests a large amount of inter-species transmission is occurring in East Cumbria, the next critical question is in what direction? We used the Bayesian Structured coalescent Approximation (BASTA) package [42] with the Bayesian evolutionary analysis platform BEAST2 (Bayesian Evolutionary analysis by Sampling Trees; [61]) to estimate *M. bovis* inter-species transmission rates. The BASTA package was able to estimate these rates whilst accounting for the known structure and sampling biases in the study population. In the current study, the sampled *M. bovis* population was split into four different sub-populations based on host species (badger or cow) and location (Cumbria or TVR) (Figure S4.1). In addition, to estimate transmission rates in a structured population, BASTA is robust to sampling biases, which are likely to be present in the current *M. bovis* dataset. Importantly, it is assumed that transmission from TVR to Cumbria only occurred in one direction, as the Cumbria clade is monophyletic within the larger TVR phylogeny (Figure 2).

The evolutionary analyses using BASTA require that there is a temporal signal in the *M. bovis* genomic data. With the presence of a temporal signal, the accumulation of substitutions will be tied to the evolutionary processes of the sampled population, making it possible to leverage genetic variation to estimate evolutionary dynamics such as transmission rates. A root-to-tip versus sampling time regression was used to determine whether a measurable temporal signal was present in the *M. bovis* genomic data (Figure S4.2). The positive trend observed in this regression indicates the presence of a weak temporal signal, therefore it was possible to proceed with the evolutionary analyses in BASTA.

The computational complexity of the analyses to be conducted using BASTA meant that the large number of *M. bovis* genomes available within the outbreak clade (orange clade in Figure 2) had to be sub-sampled.

In addition, it was only from 2014 to 2017 that the sampling of cattle and badgers in the TVR area can be considered approximately equal (in terms of effort). Therefore, only genomes sourced from infected cattle and badgers sampled from 2014 to 2018 were included in the BASTA analyses. The window was extended to 2018 to include the genomes sourced from Cumbria. The sub-sampling was then weighted to include samples from as many years as possible and from equal numbers of cattle and badgers. The sub-sampling was conducted 10 times, each time selecting 20 badgers and 20 cattle-derived *M. bovis* genomes from Cumbria and 40 badger and 40 cattle-derived genomes from the TVR area (an example of a sub-sample is shown in Figure S4.3). A root-to-tip versus sampling time regression was conducted for each sub-sample and found to be positive in all cases. Each sub-sample was then analysed separately in BASTA.

