## Supplementary material for "Phylodynamic analysis of an emergent *Mycobacterium bovis* outbreak in an area with no previously known wildlife infections": Figure S

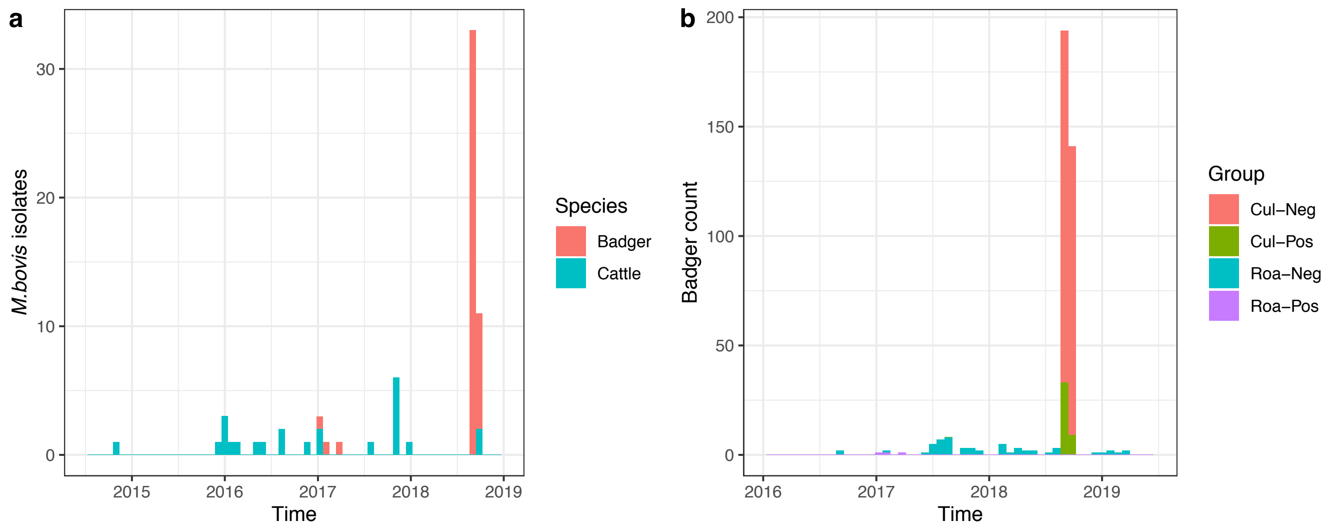


**Figure S1.1. East-Cumbria outbreak *M.bovis* isolates and badger sampling timelines.** a) The number of *M.bovis* (genotype 17:z) isolates reported in the East Cumbria outbreak, divided by animal species (farm for cattle, and badger). b) The number of badger carcasses inspected from 2016 to mid-2019 in East Cumbria, including information about their TB status (positive, Pos, or negative/uncertain, Neg) and whether they were road-kills (Roa) or trapped during the 2018 badger cull operations (Cul).

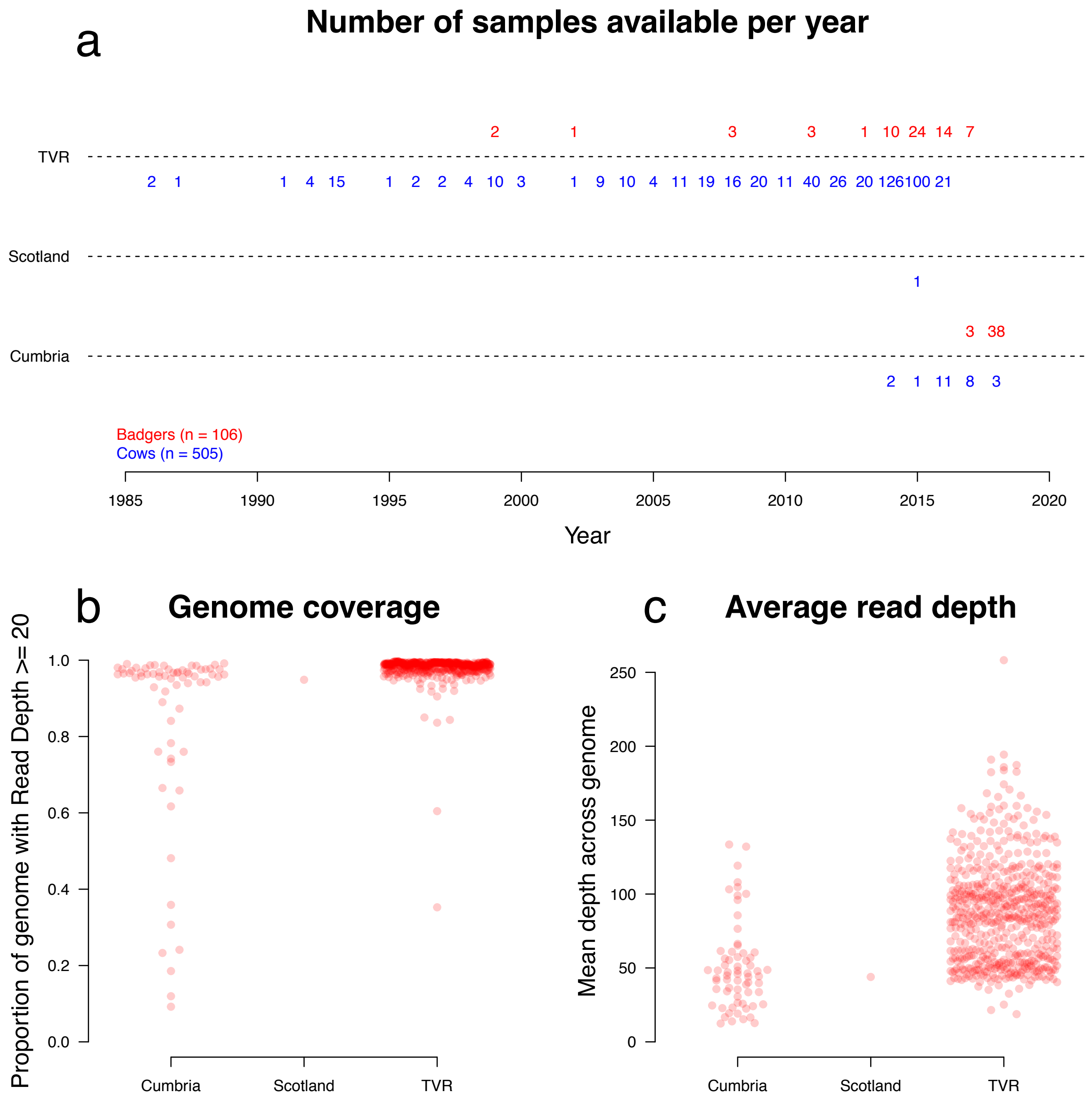


**Figure S1.2. East-Cumbria and TVR timeline and sequencing quality.** a) The sampling times of the *M. bovis* associated with the Cumbria outbreak and the Test, Vaccinate, or Remove (TVR) trial in Northern Ireland; b) the proportion of each genome sites that had >= 20 sequencing reads aligned to them for each isolate, and c) the average depth across the genome for each sequence. The cattle sampled in Scotland was eliminated from the analysis, as it was shipped for slaughtering purposes only.

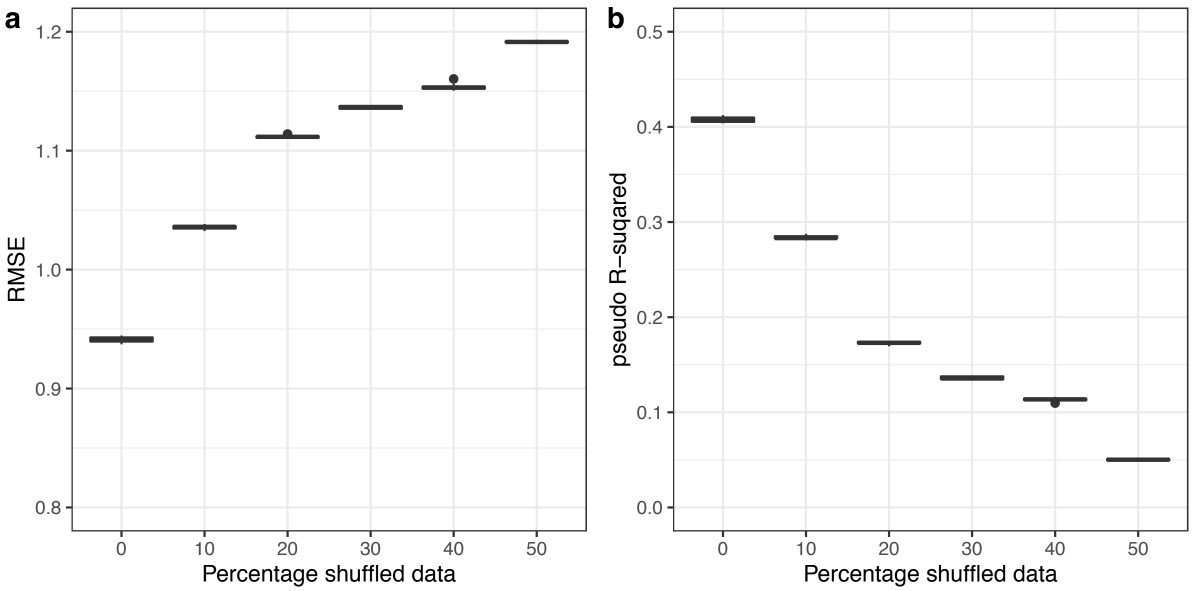


**Figure S2.1. Data shuffling results.** Effect of shuffling part of the data (from 0 to 50%) on the root mean squared error (RMSE) and the pseudo $R^2$ of the BRT model applied to the whole population (including cattle, badger, and interspecies SNV distances). The shuffling has been done by randomly reassigning a percentage of the SNV observed distances to the other covariates.

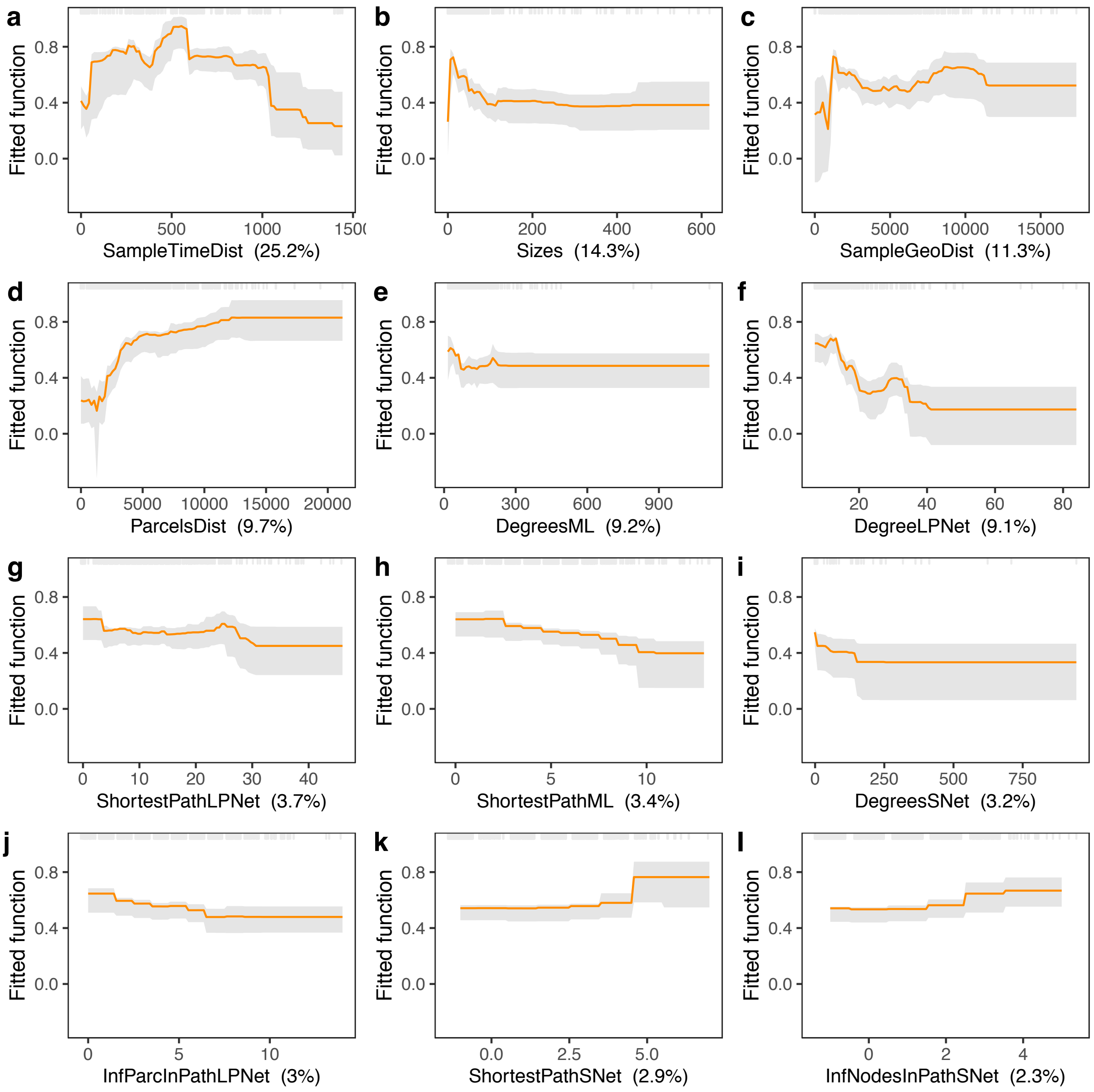


**Figure S2.2. Partial dependency plots for the BRT model.** Effect of the twelve most important epidemiological variables on the SNV distance, calculated by the boosted regression trees (BRT) model (applied to the whole population, including cattle, badger, and interspecies SNV distances).

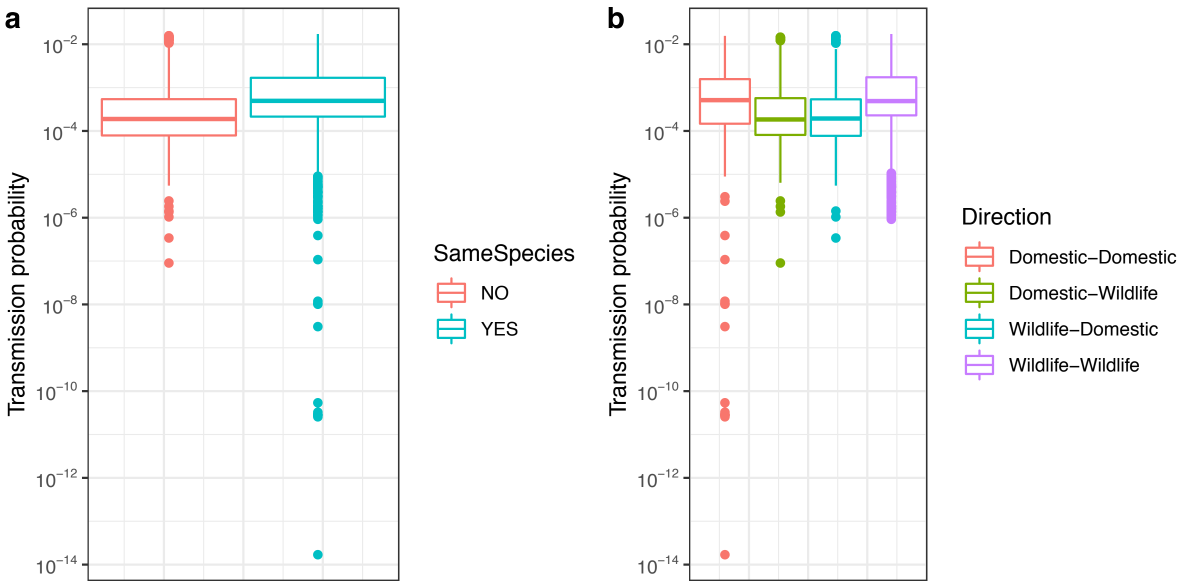


**Figure S3.1. Transmission probability by group.** Transmission probability (calculated with the Kolmogorov Forward Equation model) against groups defined by species similarity (panel a) and by transmission direction (b).

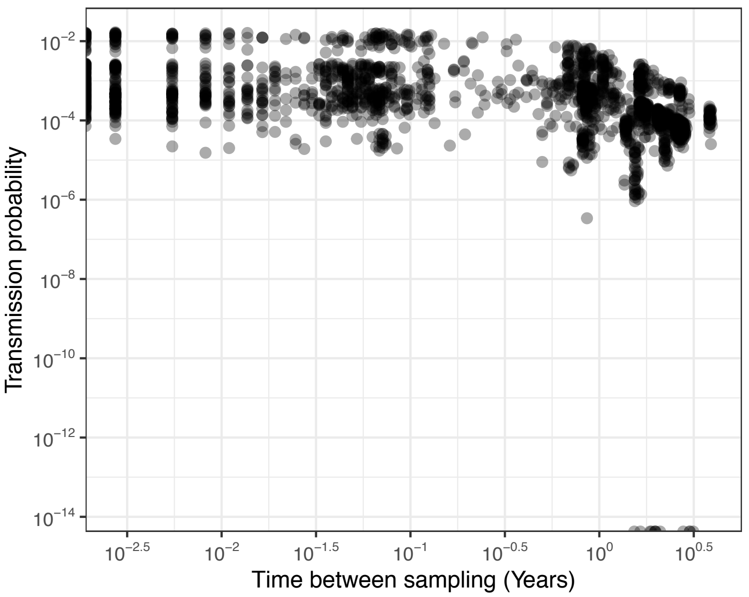


**Figure S3.2. Transmission probability vs. time.** Transmission probability (calculated with the Kolmogorov Forward Equation model) plotted against the sampling time difference between pairs'.

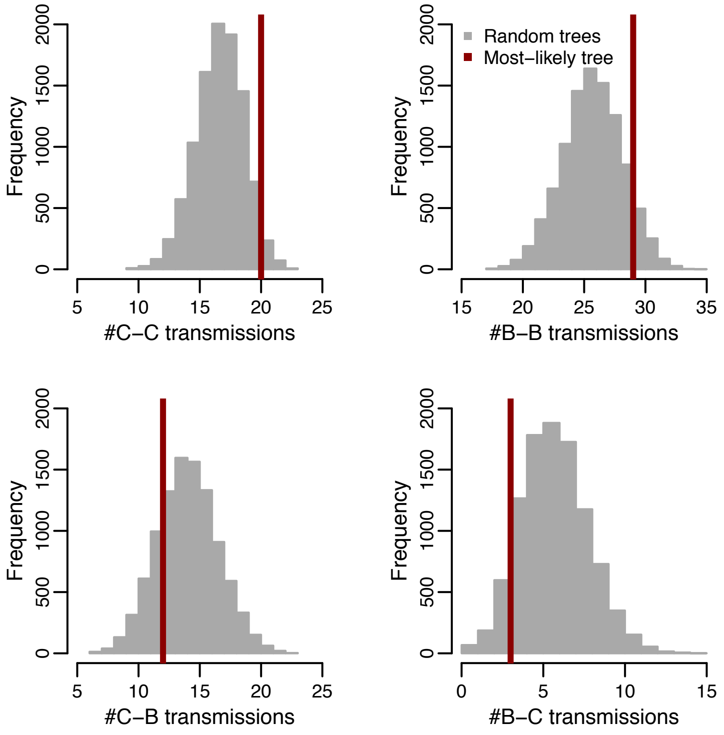


**Figure S3.3. Transmission direction in most likely vs. random trees.** Distributions of the number of transmissions divided by direction (C-C: cattle-to-cattle, B-B: badger-to-badger, C-B: cattle-to-badger, and B-C: badger-to-cattle) obtained in 10,000 random trees (grey) compared to the number of transmissions in the most likely tree (red line).

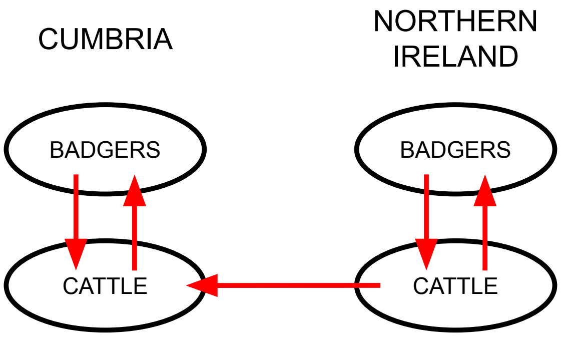


**Figure S4.1. The population structure defined for the BASTA analyses.** The red arrows indicate the different directions that *M. bovis* was permitted to transition between the different populations. TVR refers to the Test, Vaccinate or Remove trial area in Northern Ireland.

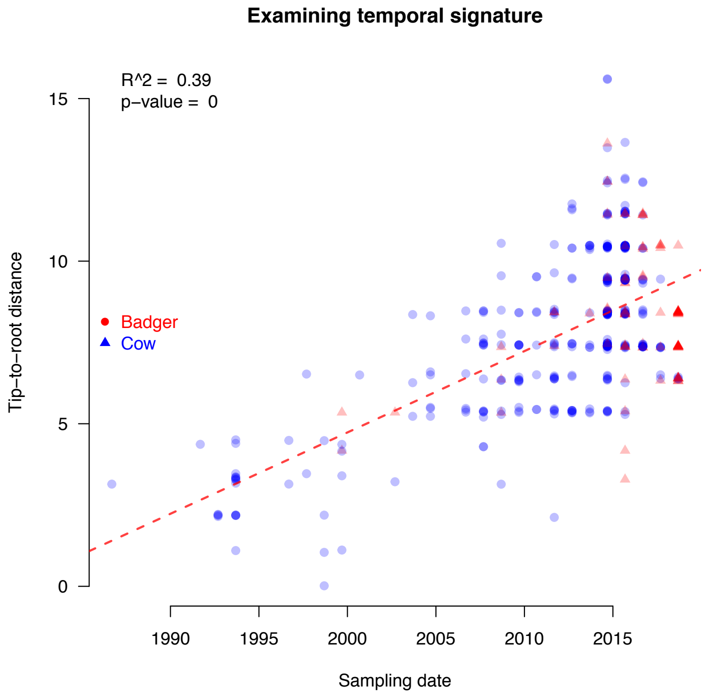


**Figure S4.2. Temporal signal in *M.bovis* isolates.** Comparing the tip-to-root distances versus the sampling time associated with each tip of the out- break clade (highlighted with orange branches in Figure 3). The $R^2$ value and red dashed line of best fit were produced by fitting a linear model in R.

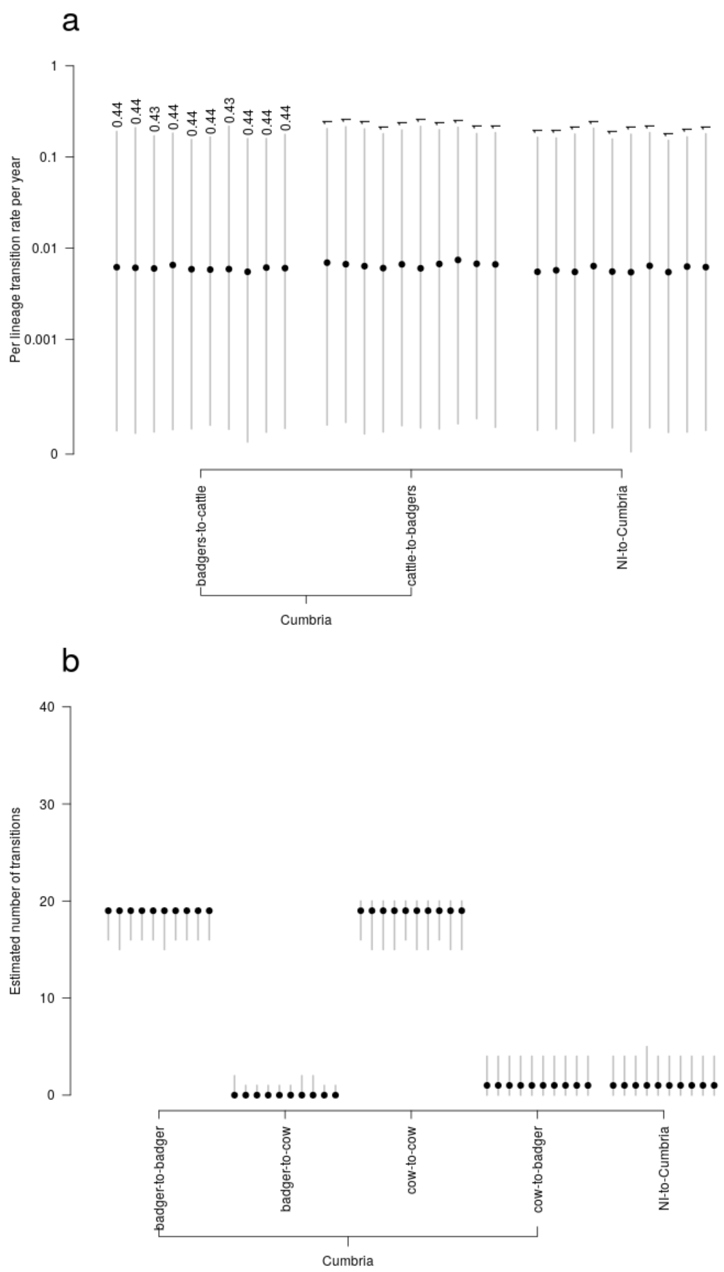


**Figure S4.3. Transmission rates and number of transmissions estimated with BASTA.** a) Inter-species and Northern Ireland to Cumbria *M. bovis* transmission rates (y axes, log scale) estimated using BASTA; b) conservative estimates from BASTA of the number of transmission events (y axes) between the sampled cattle and badgers. Conversely to the results reported in Figure 7 of the main text, here we neglected the genomic information, to test for the information present in those.

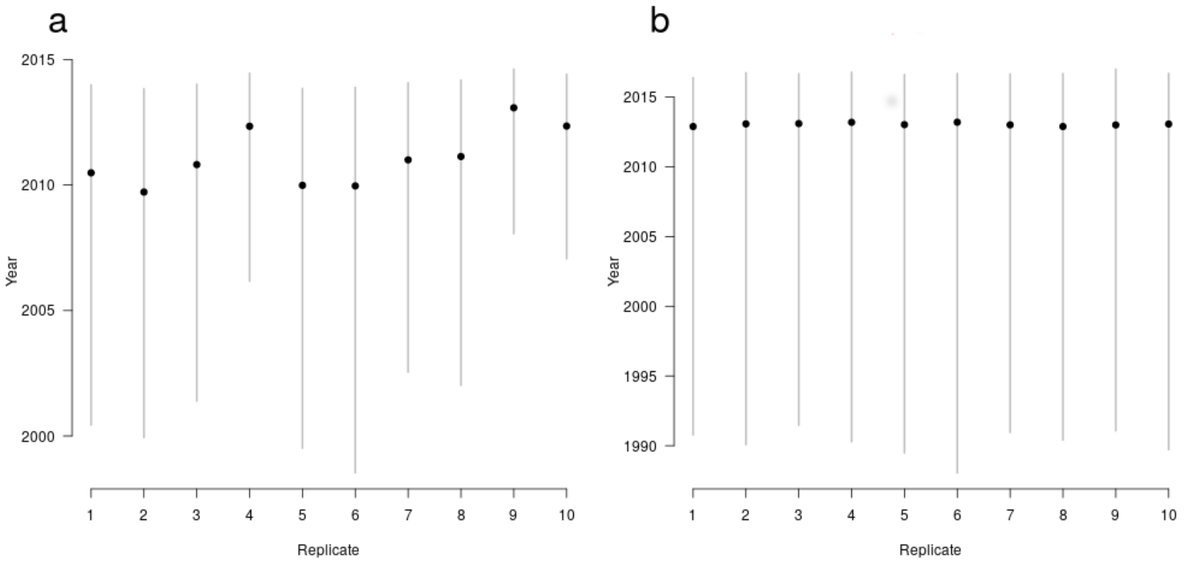


**Figure S4.4. Infected movement from TVR to East Cumbria estimate.** BASTA's estimated date for the transmission event from cattle in the TVR (Test, Vaccinate or Remove) area to cattle in Cumbria. a) The estimates derived from analyses using the genomic data. b) Estimates derived from analyses without the genomic data. In both panels the date estimates are derived from BASTA analyses of 10 different sub-samples of the available *M.bovis* genomic data.
